# Age-Related Cerebral Ventriculomegaly occurs in Patients with Primary Ciliary Dyskinesia

**DOI:** 10.1101/2024.08.12.607515

**Authors:** Franziska Eisenhuth, Joy E. Agbonze, Adam M.R. Groh, David A. Rudko, Jo Anne Stratton, Adam J. Shapiro

## Abstract

Primary ciliary dyskinesia (PCD) is a genetic disorder causing motile ciliary dysfunction primarily affecting the respiratory and reproductive systems. However, the impact of PCD on the central nervous system, through dysfunction of motile cilia in multiciliated ependymal cells, remains poorly understood. We hypothesized that patients with PCD exhibit sub-clinical ventriculomegaly due to ependymal ciliary dysfunction, which may influence neuropsychiatric diagnoses.

We demonstrated highly specific expression levels of known PCD-related genes in human brain ependymal cells (*p*<0.0001), supporting their potential role in regulating ependymal ciliary function. Computed tomography sinus images from patients with PCD (*n*=33) and age/sex-matched controls (*n*=64) were analysed. Patients with PCD displayed significantly larger ventricular areas (*p*<0.0001) and Evans index (*p*<0.01), indicating ventriculomegaly that was consistent across all genetic subgroups. Ventricular enlargement correlated positively with increasing age in patients with PCD compared to controls (*p*<0.001). Chart review demonstrated a high prevalence (39%) of neuropsychiatric/neurological disorders in adult PCD patients that did not correlate with degree of ventriculomegaly.

Our findings suggest that patients with PCD may have unrecognized, mild ventriculomegaly potentially due to ependymal ciliary dysfunction which correlates with ageing. Further study is required to determine if ventricular enlargement contributes to neuropsychiatric/neurological or other morbidity in PCD.

## Introduction

Primary ciliary dyskinesia (PCD) is a rare genetic disorder affecting motile ciliary function, with a prevalence of at least 1:7600 worldwide.^1^ In the respiratory tract, approximately 200 motile cilia extend from the apical surface of each epithelial cell and have a characteristic “9+2” ultrastructure including a ring of nine peripheral microtubule doublets surrounding a microtubule central pair. On the peripheral microtubules, outer and inner dynein arms provide motor function for ciliary beating, while the central pair, radial spokes, and nexin-dynein regulatory complexes assist with ciliary axoneme formation and beat stability. PCD is caused by variants in >50 unique genes encoding proteins in any one of these structural components, resulting in outer dynein arm (ODA) defects, outer dynein arm plus inner dynein arm (IDA) defects, IDA defects plus disorganization of the “9+2” microtubule structure, central apparatus/radial spoke (CA/RSP) defects, or oligociliary defects with vastly reduced ciliary numbers per epithelial cell.^2^

Normally, respiratory cilia beat in a coordinated, linear pattern to clear inhaled antigens trapped in mucus from the respiratory tract. In PCD, ciliary dysfunction affects respiratory cilia in the upper and lower airways, nodal cilia in the left-right organizer of developing embryos, and reproductive cilia/flagellae in sperm tails and in fimbriae of the fallopian tubes. As a result, people with PCD experience a variety of symptoms including chronic productive cough from infancy, recurrent pneumonia/bronchitis with development of bronchiectasis, chronic sinusitis, recurrent otitis media, left-right organ laterality defects (situs inversus totalis or situs ambiguous), and male and female subfertility. While the functional implications of PCD mutations in the respiratory, reproductive, and embryologic systems are well-described, the effects of ciliary mutations on central nervous system health are less understood in people with PCD.

The cerebral ventricles are lined with multiciliated ependymal cells where motile cilia on these cells beat in synchrony to aid in local circulation of cerebrospinal fluid (CSF).^3^ A preponderance of murine PCD models suggests that disruption of ependymal ciliary function leads to hydrocephalus, a condition characterized by excessive accumulation of CSF in the cerebral ventricles. In knock out mouse models of PCD, variants in genes encoding most ciliary structural components affect ependymal ciliary beating and appear to cause congenital hydrocephalus leading to early animal mortality.^4^ However, specific PCD-related genotypes initially discovered as hydrocephalus causing defects in mice (e.g. *HYDIN*, resulting in central apparatus defects) have not been similarly associated with hydrocephalus in humans.^5^ Extensive phenotyping of large PCD patient populations show that most humans with PCD do not develop hydrocephalus, aside from those with rare variants in oligociliary genes (*FOXJ1, MCIDAS, CCNO*).^6,7,8^ Among these oligociliary gene variants, there is considerable phenotypic diversity in human disease. All PCD cases with *FOXJ1* variants develop hydrocephalus, while only a minority of patients with PCD from *MCIDAS* or *CCNO* variants display hydrocephalus. The cause for this heterogeneity in humans, as well as the discrepancy between human and rodent studies, is not clear and is a topic of rigorous debate.^4^

With the pervasiveness of ependymal ciliary dysfunction and resultant hydrocephalus in most murine PCD models, we hypothesize that people with various genetic forms of PCD may have unrecognized, subtle ventriculomegaly due to ependymal ciliary dysfunction. While their ventricular enlargement may not present with clinical symptoms severe enough to warrant neurologic evaluation or brain imaging, most patients with PCD undergo computed tomography (CT) of the head to evaluate chronic sinus disease, providing a means to indirectly assess their intracranial anatomy. We further hypothesize that mild ependymal ciliary dysfunction and ventriculomegaly in people with PCD may contribute to neuropsychiatric/neurologic morbidity.^9^ Here, we demonstrate that PCD-associated genes are exclusively expressed by human ependymal cells in the brain and provide the first evidence to suggest that there is mild ventricular enlargement in patients with PCD. We also demonstrate a high prevalence (39%) of neuropsychiatric/neurological disorders in adult PCD patients.

## Materials and methods

### Single cell RNA sequencing and bioinformatic analysis

Ethical approval and consent were obtained from the Montreal Neurological Institute Research Ethics Board to utilize human brain samples from patients without PCD who underwent medical assistance in dying. Periventricular tissue samples (*n*=7) were dissociated, and cell suspensions loaded into a 10x Chromium machine for GEM generation. Libraries were prepared as per manufacturer’s recommendations and subsequently sequenced. Fastq files were generated and used as input into the cellranger software, which generated filtered count matrices that were imported in R (v.4.2.3) using the Seurat (v4.3.0.1) package. Datasets were preprocessed, normalized and integrated using standard Seurat functions. PCA, UMAP and gene expression calculations were also conducted using Seurat. To determine the significance of average expression values for each PCD gene in ependymal cells compared to all other brain cell types, non-parametric Kruskal-Wallis tests and Dunn’s multiple comparison’s tests were conducted in GraphPad Prism (v8.0.1) using raw counts that were exported from R.

### CT image analysis

#### Image selection

With approval from the McGill University Health Centre Research Ethics Board, we retrospectively obtained CT sinus/head images from past clinical testing (mainly for chronic sinusitis) in patients with PCD. A diagnosis of PCD was confirmed through biallelic variants within one known PCD gene and/or a disease-causing ultrastructural defect on ciliary transmission electron microscopy. For each PCD case, two unique control scans were obtained from sex and age-matched (within 6 months) trauma patients who lacked radiologic head injuries. All images were de-identified.

#### Data collection

We compared scans from PCD cases to controls on the basis of two measurements calculated using Bee DICOM Viewer software: the normalized ventricular area and the Evans index (Fig. 1A-D).^1^ The normalized ventricular area was calculated as the ratio of the largest axial area of the lateral ventricles and the head circumference. The Evans index, a validated clinical measurement used to diagnose normal pressure hydrocephalus, was calculated as the ratio of the maximum width of the frontal horns of the lateral ventricles and the maximal internal diameter of the skull at the same level in axial CT.^2^ Both measures account for variability in head size between individuals since the ratios are normalized by head circumference and internal diameter of the skull, respectively. Finally, clinical charts of PCD cases were reviewed for the presence of neurologic or psychiatric illness.

**Figure 1.**
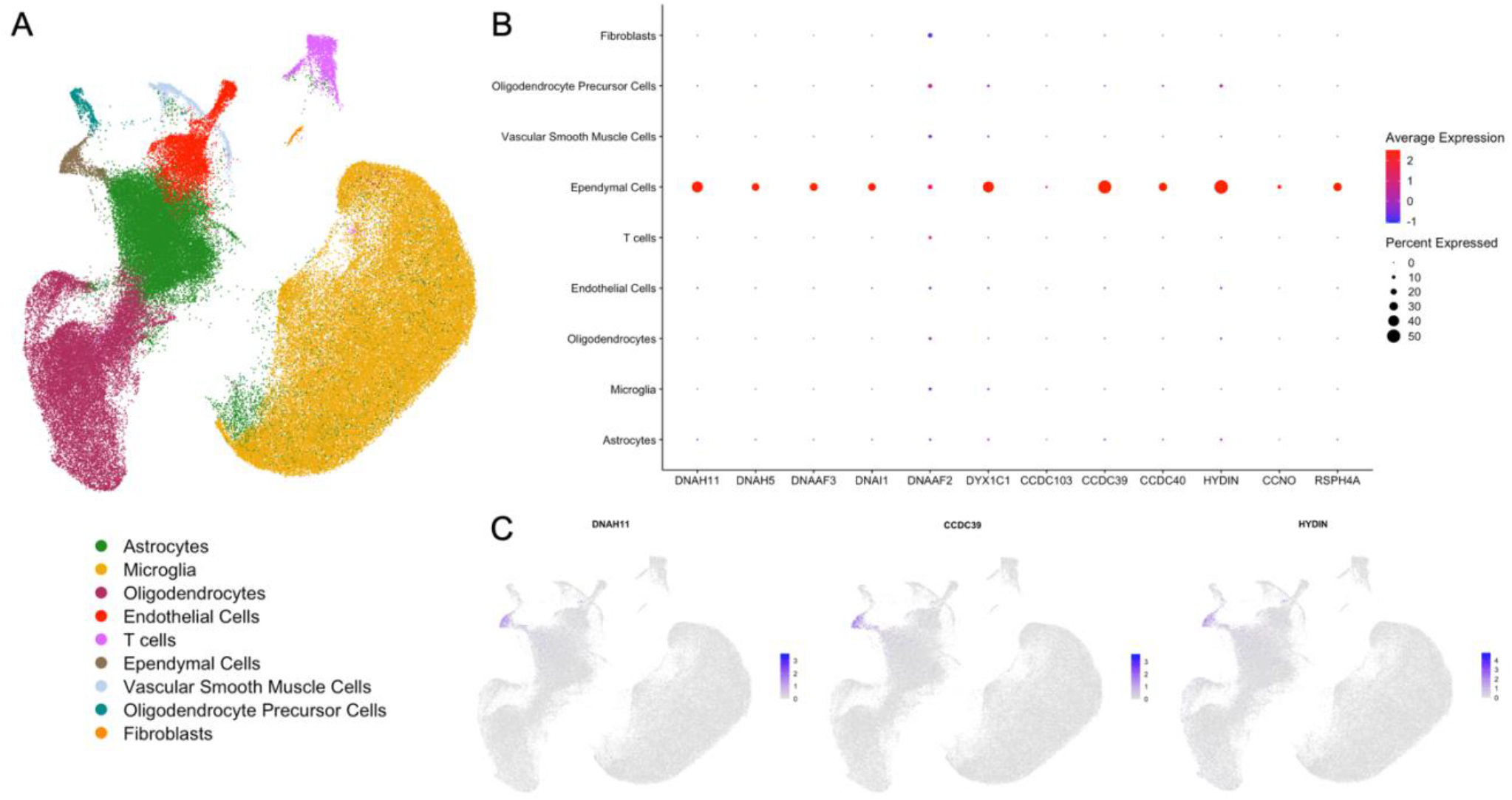
PCD-associated genes are specifically expressed in brain ependymal cells. **(A)** UMAP of cell types captured from sequencing of 13 periventricular human datasets from 7 patients. The ependymal cell cluster is identified by a brown colour. **(B)** Dot plot of PCD gene expression in all periventricular cell types (fibroblasts, oligodendrocyte precursor cells, vascular smooth muscle cells, ependymal cells, T cells, endothelial cells, oligodendrocytes, microglia and astrocytes) indicating that PCD genes are specifically expressed in ependymal cells (*p*<0.0001). Average expression values are denoted by a colour scale, with blue indicating low expression and red indicating high expression. Dot size is representative of the percent of cells expressing the gene. **(C)** Feature plots demonstrating specific expression of three representative PCD-associated genes in ependymal cells: *DNAH11, CCDC39* and *HYDIN*. Colour indicates gene expression level, with grey indicating low expression and blue indicating high expression.

#### Statistical analysis

Normalized ventricular areas and Evans index measurements were compared between PCD cases and control scans, and PCD cases were also grouped according to class of genetic defect for subgroup analysis: (1) IDA/MTD defects, (2) Dynein arm defects, or (3) Oligocilia/CA/RSP defects. Rank-sum and chi-square/Fisher exact tests compared normalized ventricular areas and Evans index measurements in PCD cases with controls and within PCD genetic subgroups. Multivariate logistic regressions of normalized ventricular area and Evans index were performed on multiple outcomes (PCD vs. control, PCD vs. control within genetic subgroups, and presence of neurologic or psychiatric diagnosis). Linear regressions of normalized ventricular area against age were performed for PCD and control groups. All models were adjusted for age and sex.

## Results

### PCD mutations are expressed specifically in ependymal cells

Expression levels of genes that are commonly mutated in PCD were evaluated in a human ventricular/periventricular single cell RNA sequencing dataset (Figure 1A) to determine whether expression was specific to brain ependymal cells. There was considerable variation for all PCD-associated genes we assessed, including in the percentage of ependymal cells and the average expression values per ependymal cell that expressed each gene (Figure 1C). Even so, PCD-associated genes were expressed specifically (*p*<0.0001) in human brain ependymal cells compared to other periventricular cell types captured in our samples (fibroblasts, oligodendrocyte precursor cells, vascular smooth muscle cells, T cells, endothelial cells, oligodendrocytes, microglia and astrocytes) (Figure 1B and C).

### Lateral ventricles are larger in PCD and increase with age

CT sinus/head scans were available for 47 patients with confirmed PCD, but only 33 patient scans captured enough of the cerebral ventricles to allow for full analysis within our protocol (mean age 27.3 ± 15.2 years, 12 pediatric <18 years old, 21 adult ≥18 years old, 12 male, 21 female). These were compared to 64 age and sex matched trauma control scan patients, who lacked any evidence of cerebral pathology on their imaging. In light of inadequate control availability for age and sex-matching, two trauma control scans were used twice. Patients with PCD showed larger mean normalized ventricular areas (PCD 0.21 ± 0.12, control 0.12 ± 0.05, *p*<0.0001) and mean Evans index measurements (PCD 0.26 ± 0.05, control 0.22 ± 0.06, *p*<0.01) compared to controls (Fig. 2A-B). Logistic regression examined the association between normalized ventricular area or Evans index and the presence or absence of PCD. The odds ratios for normalized ventricular area and Evans index were 1.202 (95% *CI*: 1.10–1.31) and 1.139 (95% *CI*: 1.04–1.25) respectively, indicating that individuals with larger cerebral ventricles were more likely to belong to the PCD group than the control group (Table 1). These associations remained statistically significant for normalized ventricular area (*p*<0.0001) and Evans index (*p*< 0.01) after adjusting for age and sex. Three patients in the PCD group had an Evans index above the cut-off (0.31) for a clinical diagnosis of ventriculomegaly. Linear regression of normalized ventricular area against age revealed that ventricular size increased markedly with age in patients with PCD (*β*=0.005, *p*<0.0005) but remained relatively constant in the control group (*β*=0.0009, *p*<0.05) (Fig. 1C).

**Figure 2.**
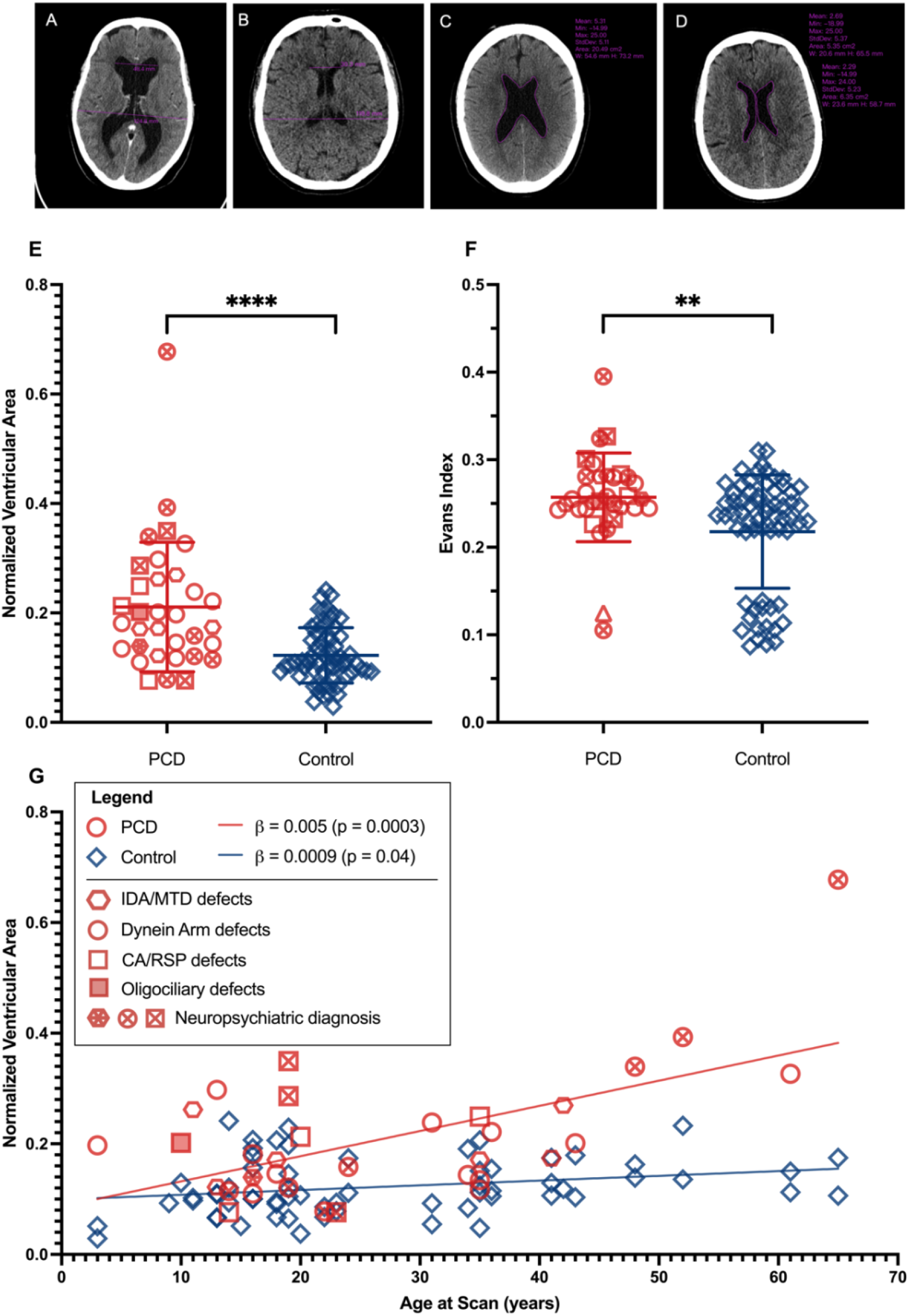
Normalized ventricular area and Evans index are larger in PCD that control groups. **(A)** Axial slice of the head CT from a patient with PCD from *DNAH5* variants with Evans index measurement - ratio of the maximum width of the frontal horns of the lateral ventricles and the maximal internal diameter of the skull at the same level. (**B**) Axial slice of head CT from corresponding age and sex matched control for the patient in (A) with Evans index measurement. (**C**) Axial slice of head CT from patient with PCD from *DNAH11* variants, with measurement of ventricular area. **(D)** Axial slice of head CT from corresponding age and sex matched control for patient in (C) with measurement of ventricular area. (**E**) Mean normalized ventricular area is significantly larger in patients with PCD (0.21±0.12) than in the Control group (0.12±0.05), \*\*\*\**p*<0.0001. (**F**) Mean Evans index is significantly larger in patients with PCD (0.26±0.05) than in in the Control group (0.22±0.06), \*\**p*<0.01. (**G**) Normalized ventricular area plotted against age shows that ventricle size increases significantly more with advancing with age in patients with PCD (*β*=0.005, *p*<0.001) than in controls (*β*=0.0009, *p*<0.05).

**Table 1.**
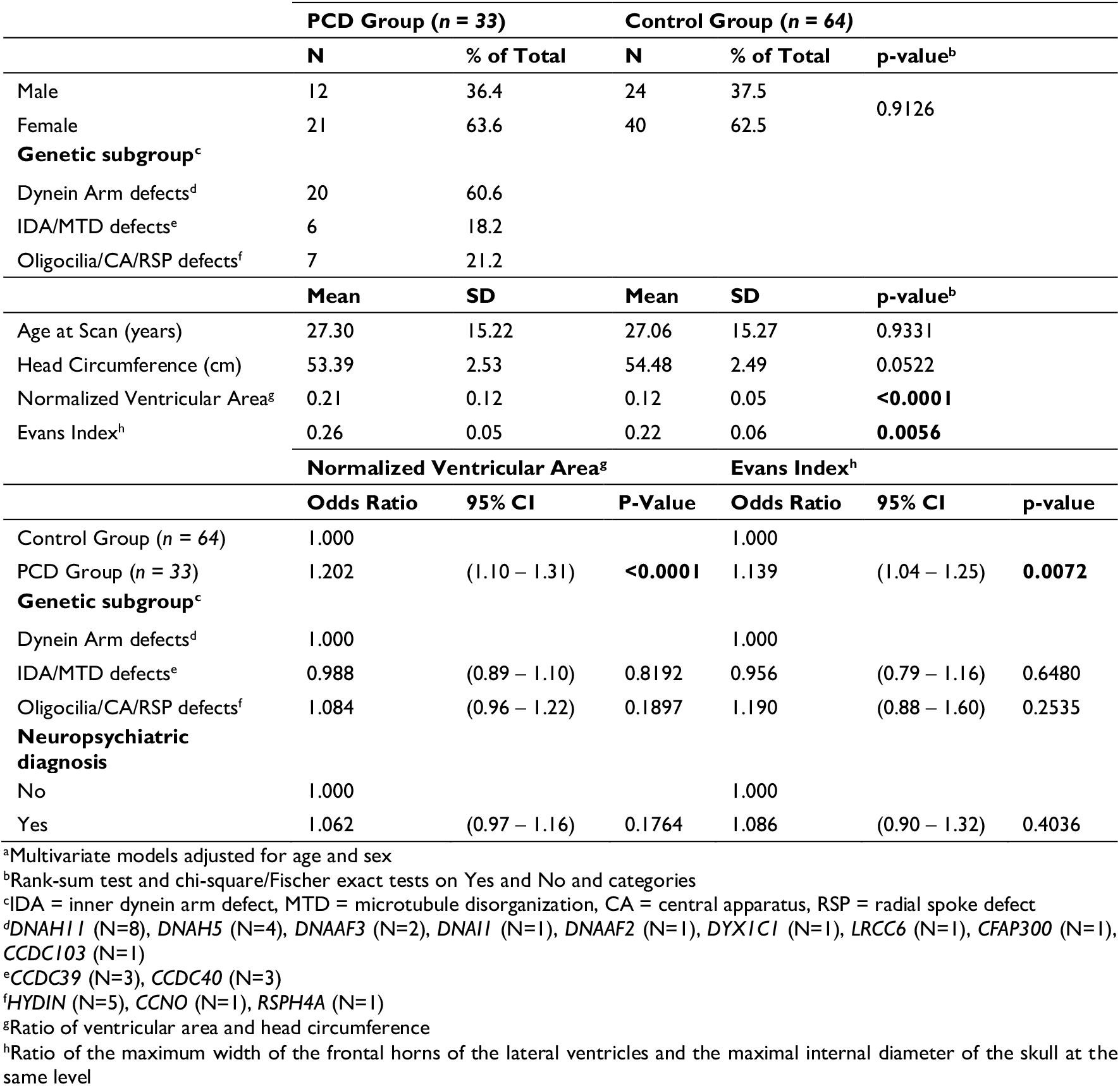
Descriptive statistics and logistic regression of normalized ventricular area and Evans index on multiple outcomes^a^.

Computed tomography scans from patients with PCD were categorized into three genetic subgroups depending on the PCD gene affected: (1) IDA/MTD defects (*n*=6 total, with *n*=3 *CCDC39, n*=3 *CCDC40*), (2) Dynein arm defects (*n*=20 total, with *n*=8 *DNAH11, n*=4 *DNAH5, n*=2 *DNAAF3, n*=1 for each of *DNAI1, DNAAF2, DYX1C1, LRCC6, CFAP300, CCDC103*), or (3) Oligocilia/CA/RSP (*n*=7 total, with *n*=1 *CCNO, n*=5 *HYDIN, n*=1 *RSPH4A*). Subgroup analysis revealed similar increases in normalized ventricular area and Evans index across all genetic subgroups when compared to controls. Logistic regressions conducted for normalized ventricular area and Evans index on genetic subgroup as an outcome showed no significant positive odds ratios for the IDA/MTD defects group (*p*=0.8 for area, *p*=0.6 for Evans) or the Oligocilia/CA/RSP group (*p*=0.2 for area, *p*=0.3 for Evans) compared to the Dynein arm defects group.

### High prevalence of neuropsychiatric/neurological illness in PCD

Medical chart reviews were performed in patients with PCD for presence of neurologic or psychiatric conditions. In our PCD cohort, a diagnosis of neurologic or psychiatric illness was present in 39% of adults (with only one patient being purely a neurologic diagnosis). Diagnoses included depression, dyslexia, schizophrenia, anxiety, migraines, leg weakness/paresthesia, and vision loss. Depression was the most common diagnosis, which was present in 36% of adults (Table 2). Pediatric patients with PCD had less of a neuropsychiatric burden, with 25% affected mainly by attention deficit/hyperactivity disorder, autism, or developmental delay. When including adult patients only with neuropsychiatric conditions, 33% of the PCD cohort had a neuropsychiatric diagnosis, compared to the 12.5% population prevalence of neuropsychiatric illness as per the World Health Organization.^10^ However, normalized ventricular area and Evans index did not correlate with the presence of a neuropsychiatric disorder (*p*=0.2 and *p*=0.4, respectively).

**Table 2.** Neurologic and psychiatric diagnoses in patients with PCD.

| Age | Sex | Genetic Subgroup<sup>a</sup> | PCD Gene | Diagnosis |
| --- | --- | --- | --- | --- |
| 14 | F | Dynein arm defect | DNAH5 | ADHD<sup>b</sup> |
| 16 | F | IDA/MTD defect | CCDC40 | ADD<sup>c</sup> |
| 19 | F | Oligocilia/CA/RSP defects | HYDIN | Depression, ADD |
| 19 | M | Dynein arm defect | LRRC6 | Autism, developmental delay |
| 19 | M | Oligocilia/CA/RSP defects | HYDIN | ADHD, dyslexia |
| 22 | F | Dynein arm defect | DNAH5 | Schizophrenia |
| 23 | F | Oligocilia/CA/RSP defects | RSPH4A | Depression, anxiety, ADHD |
| 24 | M | Dynein arm defect | DNAAF2 | Migraines, depression |
| 48 | F | Dynein arm defect | DNAAF3 | Depression |
| 52 | F | Dynein arm defect | DNAH11 | Depression |
| 65 | F | Dynein arm defect | DNAH5 | Leg weakness/paresthesia, vision loss |

<sup>a</sup>IDA = inner dynein arm defect, MTD = microtubule disorganization, CA = central apparatus RSP = radial spoke defect
<sup>b</sup>Attention deficit hyperactivity disorder
<sup>c</sup>Attended deficit disorder

## Discussion

PCD is a genetic disorder which impairs motile ciliary function throughout the body. While the effects of motile ciliary dysfunction are well described in the human respiratory and reproductive tracts, pathogenesis related to motile cilia defects in ependymal cells of the central nervous system remains unclear. Most PCD-related gene mutations cause severe hydrocephalus in murine models, yet in humans, hydrocephalus only arises with variants in a few, rare PCD genotypes causing oligociliary defects. Our analysis clarifies that in fact people with PCD *do*, more generally, have abnormal features of the brain consistent with CNS-driven motile ciliary dysfunction. Several patients with PCD even had frank ventriculomegaly with Evans indices above the diagnostic cutoff value of 0.31.^11^ Based on our gene expression data, PCD genes were highly, and largely specifically expressed by ependymal cells, suggesting defects in this glia cell type might be a driver of ventriculomegaly. Though most patients with PCD usually do not develop clinically appreciable hydrocephalus, we propose that ependymal cell dysfunction in PCD may lead to sub-clinical ventriculomegaly. This subtle enlargement of the cerebral ventricles may have implications for the development of neurologic and psychiatric illness in PCD patients.

Ventricular enlargement in PCD seems to worsen with age, as adults >40 years old generally showed larger differences in normalized ventricular area from age-matched controls. Given this, we were curious whether this had any relation to idiopathic normal pressure hydrocephalus (iNPH), which is also a disease characterised by ventriculomegaly where prevalence increases with age. There are some suggestions of cilia-related defects in genome-wide association studies of iNPH.^12^ Studies have recently implicated *CFAP43* and *CWH43* as genes which drive ciliary abnormalities that were associated with iNPH.^13,14^ However, no epidemiology studies have suggested an association between PCD and iNPH as of yet.^15^ Our data includes mostly young patients with PCD (<40 years old), which is a common problem with PCD cohorts (a relatively young research field). Diagnosis of PCD is generally established during childhood, with an average age of diagnosis near 6 years.^16^ Adults with bronchiectasis and chronic respiratory disease are rarely considered for PCD testing, as their disease is usually blamed on other, more common conditions, like asthma or chronic obstructive pulmonary disease. As a result, most PCD research cohorts worldwide are skewed to include younger patients, and with this age bias, researchers cannot easily appreciate the ventriculomegaly that appears to worsen in older patients. Though PCD is generally associated with a normal life expectancy, very few cases are published in patients older than 60 years of age, which may also underlie the lack of age-associated neurologic observations in this rare disease. To date, there are no publications broadly linking PCD with neuropsychiatric or neurologic disease.

iNPH is classically associated with gait disturbance, cognitive impairment, and urinary incontinence in patients older than 60 years of age. Though none of our patients displayed these classic symptoms, many non-specific neurologic issues and psychiatric manifestations have been linked to ventriculomegaly and hydrocephalus.^17,18,19^ It is unclear if the observed increase in neuropsychiatric illness in our PCD cohort was due to psychosocial stress stemming from having a chronic respiratory disease or due to their ventriculomegaly. One recent analysis of psychological screening in patients with PCD showed 30-40% likely had depression or anxiety.^20^ Further study of central nervous system issues in larger PCD patient cohorts, including in-depth neuropsychiatric testing/questionnaires and longitudinal functional brain imaging, may help to determine if ventricle size directly influences neuropsychiatric outcomes.

Genotype-phenotype correlation studies in PCD have consistently shown that IDA/MTD defects or biallelic variants in corresponding *CCDC39* or *CCDC40* genes result in severe respiratory and nutritional outcomes compared to PCD caused by dynein arm defects.^21,22^ This trend was not seen in the ventricular effects of our PCD cohort, as the three genetic subtype groups, including IDA/MTD defects, have similar measures of ventricle size. In other PCD genotypes affected by splice variants or hypomorphic changes, respiratory outcomes may be milder.^23,24^ However, our relatively small number of PCD cases did not afford enough power to perform this type of genetic analysis. Perhaps with larger cohorts of PCD patients, including a wide spectrum of genotypes, genotype-phenotype relationships would be apparent.

The discrepancies between ventriculomegaly in humans and rodent models is a current topic of debate, so our findings are timely.^25^ Many hypothesize that the reason PCD variants have such a devastating impact in smaller animals is that ependymal cilia motility is more important in narrower tubes and ventricular systems due to their small size. The fact that ependymal motile cilia do not scale with ventricle size in higher-order animals lends support for this notion. However, multiple PCD genotypes in our cohort contributed to the overall effect of ventriculomegaly in patients with PCD, suggesting that ependymal cilia, however small, do still play an important role in maintaining ventricular homeostasis, even in humans.

Interestingly, ventriculomegaly has been reported on prenatal ultrasound in a small number of infants diagnosed with PCD, however their ventriculomegaly *apparently* resolves by the time of birth.^26^ The ependymal cell layer begins to differentiate in the human brain as early as 21 weeks gestational age (and fully matures by ∼ 7 months of age), so the observed prenatal ventriculomegaly may be a direct result of the absence of functional motile cilia in early embryonic development.^27^ However, this does not explain the age-related increase in ventriculomegaly that we observed in adult patients with PCD. Our finding that ventriculomegaly becomes more severe with age could suggest that vasomotor tone and gravity (the main drivers of bulk CSF flow) are able to partially compensate for reduced ependymal ciliary beating for most of one’s life. The reason for this is unknown but could be due to the accumulation of toxic solutes over time and alterations in local CSF exchange with interstitial fluid.^28,29,30^

In conclusion, our results implicate PCD in a neurologic/neuropsychiatric disease process on top of its well-studied effects on the respiratory and reproductive systems. We found that patients with PCD have sub-clinical ventriculomegaly which worsens with age and is likely caused by ependymal ciliary dysfunction. Given the high prevalence of neuropsychiatric illness in this population, neurologic and psychiatric screening and evaluation should be considered in patients with PCD.

## Data availability

The data that support the findings of this study are found in the supplementary files.

## Acknowledgements

A kind thank you to the patients with PCD whose data was used for this study; to Frederic Lamonde, Raman Agnihotram, Biostatistics Consulting Unit of the McGill University Health Center; to Debbie Friedman and Tarek Razek, Trauma program at the McGill University Health Centre. This work was presented as an abstract at the American Thoracic Society Conference in 2024, San Diego, California, USA.

## Supplementary material

Supplementary material is available at *Brain* online.

## Footnotes

1 Sainuo United Medical Technology (Beijing) Co., Ltd. Bee DICOM Viewer. Version 2.4.3. Apple Store; 2020. [Software].

2 Relkin N, Marmarou A, Klinge P, Bergsneider M, Black PM. Diagnosing idiopathic normal-pressure hydrocephalus. Neurosurgery 2005; 57(3 Suppl.): S4–S16; discussion ii–v.

